# Uncovering Carbohydrate Metabolism and Endogenous Hormone Regulation during Flush Phenology in Citrus Trees using Proteomics and Metabolomics

**DOI:** 10.64898/2026.03.09.710605

**Authors:** Kuikui Chen, Syed Bilal Hussain, Xin Liu, Qingyan Meng, Christopher Vincent, Yu Wang

## Abstract

Rapid shoot growth (flushing) phenology is a fundamental developmental process in perennial woody plants such as citrus. In a separate study, we identified physiological shifts from photosynthesis to mobilization of nitrogen and carbohydrate to support new shoot growth. However, the underlying molecular and biochemical signals remain largely unknown. Here, we integrated proteomic and metabolomic analyses to investigate carbohydrate and hormone dynamics across three flush stages in *Citrus sinensis*: quiescent period (stage 1), new shoot initiation (stage 2), and full expansion (stage 3). Sucrose, maltose, and trehalose accumulated in apical leaves during early shoot initiation and declined during subsequent shoot expansion, indicating depletion of carbohydrate reserves and enhanced resource remobilization. These changes were accompanied by coordinated regulation of starch-metabolizing enzymes, including ADP-glucose pyrophosphorylase, α-amylase, and isoamylase, supporting a transition from carbon storage to carbon export during active shoot growth. Indole-3-acetic acid increased continuously across stages, while trans-zeatin and gibberellin A□ showed opposite trends in apical versus basal leaves before jointly increasing at stage 3. Hormone analysis revealed dynamic and coordinated signaling changes during flush development. Abscisic acid declined from stage 1 to 2, whereas jasmonoyl-isoleucine and salicylic acid increased from stage 2 to 3. Some hormone-responsive proteins, including Gretchen Hagen 3 and Gibberellin-insensitive dwarfing 1, exhibited expression patterns consistent with hormonal fluctuations. Together, these results support a stage-specific regulatory framework in which carbohydrate metabolism and hormone signaling are tightly coordinated to regulate rapid source–sink transitions during citrus flush development.

**Highlight:** We reveal how carbohydrate metabolism and hormone signaling are spatiotemporally coordinated during citrus shoot growth phenology, and we develop an integrated metabolic–hormonal model that connects carbon allocation to developmental transitions.

## Introduction

Citrus trees (*Citrus spp.*) are economically significant fruit crops, and, like many woody species, they exhibit a flushing patter of growth (Li et al., 2020). Flush phenology consists of recurring periods of quiescence in mature shoots, followed by rapid expansion of new shoots (flushes). Under optimal conditions, the cycle repeats approximately every 6-8 weeks. Flush dynamics strongly influence vegetative growth, fruit yield, stress tolerance, and fruit quality, making it a critical component of orchard management in subtropical tree crops such as citrus and avocado (Salazar-Garcia et al., 2006). Flushing is governed by both environmental cues (light, temperature, and water availability) and internal factors, notably carbohydrate metabolism and endogenous hormones (Carvalho et al., 2021; Li et al., 2022).

Carbohydrates are central to energy supply and structural integrity during plant growth. During citrus flushing, starch degradation and carbohydrate redistribution provide essential energy for new shoot and leaf development, which must occur rapidly, as new shoots complete expansion within approximately 3 weeks after growth initiation (Muhammad et al., 2020). Starch, a primary carbon reserve, is hydrolyzed into soluble sugars such as sucrose and glucose, which supply energy for morphogenesis and metabolic processes, including leaf and floral bud development (Li et al., 2014; Bhatla et al., 2023).

Investigating carbohydrate dynamics during flushing offers insights into energy allocation strategies during this crucial phase. Understanding how trees balance the C and N demands of new growth with maintenance of the function of existing foliage could potentiate new approaches to improving agricultural production. Vincent et al. (2025) showed that mature leaves increase carbon export and deplete nearly all starch reserves to support new shoot growth, whereas Hussain et al. (*under review*) reported that photosynthesis decreases in the export phase, rather than increasing to support greater carbohydrate demand. Understanding the signaling mechanisms underlying these apparent trade-offs would clarify the balance between current photosynthesis and new growth and may facilitate more efficient transitions through flushing cycles.

In addition to their developmental roles, plant hormones are important regulators of carbohydrate metabolism. Endogenous hormones further modulate the flushing by regulating cell division, expansion, and dormancy (Cai et al., 2018; Gill et al., 2023). Auxin gradients orchestrate organogenesis and pattern formation by regulating both cell division and expansion (Perrot-Rechenmann et al., 2010). Gibberellins (GAs) primarily promote elongation growth, and exogenous GA application typically enhances the size of citrus trees (Vashisth et al., 2024). In contrast, abscisic acid (ABA) induces bud dormancy (Luo et al., 2019). Endogenous phytohormones not only regulate shoot development but also modulate carbohydrate metabolism to coordinate growth with energy supply. Auxins and cytokinins influence sucrose allocation by regulating sugar transporters and phloem loading (Sairanen et al., 2012; Roitsch & Ehneß, 2000; Ruan et al., 2010). Jasmonates (JAs) redirect carbohydrate fluxes toward defense under stress conditions (Babst et al., 2005; Gómez et al., 2010). Despite their importance, studies examining the dynamic interactions between carbohydrate metabolism and hormone signaling during citrus flushing remain limited, particularly at the molecular and metabolic levels.

Proteomics and metabolomics are powerful tools in omics research, providing complementary insights into plant physiology (Yan et al., 2022). Proteomics identifies and quantifies proteins to reveal biochemical changes, while metabolomics profiles small-molecule metabolites involved in metabolic pathways (Liu et al., 2024). Integrating these approaches enables a comprehensive understanding of enzyme and metabolite fluctuations during key physiological transitions or phenotypic changes (Li et al., 2023). In previous work, we applied proteomics and/or metabolomics to identify early chemical markers of citrus infected with huanglongbing, investigate the molecular basis of off-flavors in sweet orange fruit, and assess how moderate shading could mitigate the effects of huanglongbing in field-grown sweet oranges by reducing leaf starch accumulation and maintaining photosynthetic efficiency (Wang et al., 2022; Suh et al., 2021; Yao et al., 2019). However, the coordinated metabolic and hormonal regulatory mechanisms underlying citrus flush development remain largely unexplored using an integrated omics framework.

In this study, we employed an integrated proteomic and metabolomic approach to investigate carbohydrate metabolism and hormone regulation during flush phenology in sweet orange (*Citrus sinensis* [L.] Osbeck). By elucidating key biochemical transitions, this work advances understanding of citrus growth regulation and source–sink dynamics and provides a basis for optimizing cultivation strategies to improve yield and fruit quality.

## Materials and methods

### Plant materials

Healthy *C. sinensis* (cv. ‘Valencia 1-14-19’) grafted onto US 942 rootstock (*C. reticulata* ‘Sunki’ x *Poncirus trifoliata* ‘Flying Dragon’) were obtained from a commercial nursery in November 2022 and cultivated in a controlled greenhouse at the Citrus Research and Education Center, University of Florida (Lake Alfred, FL, USA; 28.1021° N, 81.7121° W). Upon arrival, plants were transplanted into 10-L containers filled with Pro-Mix BX potting substrate (Premier Tech Ltd., Quebec, Canada). To ensure uniform growth, all branches were pruned in February 2023. New shoots emerged within 7–10 days, and shoot numbers were standardized to 4–5 evenly distributed branches per plant. From each plant, a branch with a nearly equal number of leaves and a length of 14-15 cm was selected, resulting in 8-9 leaves on mature flushes. Greenhouse conditions were maintained at 24–35 □ with 64–76% relative humidity. Temperature fluctuations were regulated by a pad-and-fan system, which automatically activated at 30 □. Plants received 2.5 g of 20-20-20 fertilizer biweekly, and pest inspections were conducted weekly. Three developmental stages were defined for sampling: (1) mature leaves at the base and the apex of mature shoots in the quiescent period with only mature shoots (stage 1, quiescent period), (2) the period of new shoot initiation (stage 2, new shoot initiation), and (3) the period of full expansion of new shoots (stage 3, full expansion). Five trees were selected for each stage.

### Chemicals and reagents

Isotopically labeled internal standards, including L-tyrosine-¹³C□, L-proline-d_3_, L-phenylalanine-¹³C□, and hippuric acid-d_5_, were purchased from Sigma-Aldrich (St. Louis, MO, USA). LC-MS grade acetonitrile, methanol, water, formic acid, and ammonium acetate were obtained from Fisher Scientific (Fair Lawn, NJ, USA).

### Sample preparation for metabolomics

Leaf metabolite profiling, including targeted and untargeted metabolomics, was performed on samples collected from two leaf positions (apical and basal) across three growth stages (Stage 1, Stage 2, and Stage 3; n = 5). Leaf samples were freeze-dried for 24 hours and finely ground. For reverse-phase chromatography (RP) analysis, 35 mg of ground leaf tissue was extracted with 0.7 mL of 20% methanol containing L-tyrosine-¹³C□ and L-phenylalanine-¹³C□ as internal standards. For hydrophilic interaction chromatography (HILIC), a separate 35 mg aliquot was extracted with 0.7 mL of 80% methanol containing hippuric acid-d□ and L-proline-2,5,5-d□ as internal standards. Extraction was carried out by vortexing for 10 min, followed by ultrasonication for 30 min. The samples were then centrifuged (12,000 g, 4 □, 15 min), and the supernatant was filtered through a 0.22-μm syringe filter. A 2-μL aliquot of the extract was injected into the LC–HRMS system. Quality control (QC) samples were prepared and extracted by mixing equal amounts of all samples.

### Experimental Conditions for Targeted Metabolomics

The LC–HRMS system consisted of a Vanquish UHPLC coupled to a Q Exactive Plus Orbitrap mass spectrometer (Thermo Fisher Scientific, Waltham, MA, USA). Equipped with a heated electrospray ionization (HESI) interface, the mass spectrometer operated in targeted-selected ion monitoring (t-SIM) mode, enabling positive- and negative-ion switching. Detailed experimental settings are provided in the **Supplementary Information.**

### Data processing for metabolomics

For targeted metabolomics, Xcalibur (Ver. 4.0, Thermo Fisher Scientific, San Jose, CA, USA) was used for instrument control and data analysis. Metabolite concentrations were quantified using peak-area ratios relative to internal standards. In untargeted metabolomics, Xcalibur (Ver. 4.0) controlled data acquisition, while Compound Discoverer (Ver. 3.1, Thermo Fisher Scientific) processed the data, including feature detection, retention time alignment, and area normalization. Metabolite annotation was performed using exact mass, molecular formula, and MS² fragmentation patterns, with database searches against mzCloud, ChemSpider, and MassList. A mass tolerance of 2 ppm was applied, and the FISh score was used to assess the quality of MS² fragmentation. Only metabolites with a relative standard deviation (RSD) below 30% in quality control (QC) samples were included in subsequent analyses. Differential metabolites across growth stages and leaf positions were identified using volcano plot analysis, with selection criteria of fold change >1.5 or <0.67 and *p*-value <0.05 (t-test). Pathway enrichment analysis was conducted using MBRole (Ver. 2.0) to identify key metabolic pathways associated with phenology and leaf position (http://csbg.cnb.csic.es/mbrole2).

### Sample preparation for Proteomics

Samples were lysed in a buffer containing 9 M urea, 5% SDS, and 50 mM TEAB, followed by sonication and centrifugation. Proteins were precipitated using pre-chilled acetone, incubated at −20 □, centrifuged, and resuspended for quantification via the BCA assay. For digestion, 50 μg of protein was reduced, alkylated, acidified, and processed using S-Trap columns. After sequential washing, proteins were digested overnight at 37 □ with Trypsin/LysC (1:10, w/w). The resulting peptides were eluted in fractions, pooled, dried, and desalted using C18 columns before further analysis.

### Experimental Conditions for Proteomics

All fractionated samples were analyzed by nano flow HPLC (Ultimate 3000, Thermo Fisher Scientific) followed by Orbitrap EclipseTM TribridTM (Thermo Fisher Scientific). Nanospray FlexTM Ion Source (Thermo Fisher Scientific) was equipped with Column Oven (PRSO-V2, Sonation) to heat up the nano C18 column (250 mm × 75 µm, 1.7 µm) for peptide separation. Detailed analytical parameters are provided in the **Supplementary Information**.

### Data processing for Proteomics

Proteome Discoverer 2.3 (PD 2.3, Thermo Fisher Scientific, San Jose, CA, USA) was used for data-dependent acquisition (DDA) spectral library construction and subsequent data-independent acquisition (DIA) analysis. Peptides identified with a false discovery rate (FDR) < 1% were included in the final spectral library. Functional annotation, including Gene Ontology (GO), Eukaryotic Orthologous Groups (KOG), and pathway enrichment analysis, was performed within the same pipeline. For DIA quantification, MSstats, employing a linear mixed-effects model, processed the data according to predefined comparison groups and performed significance testing. Differential protein analysis was conducted using fold-change criteria of>1.5 or <0.67, with a *p*-value <0.05. Function enrichment analysis was subsequently carried out to identify key biological processes associated with differential proteins.

## Results

### Metabolomics analysis

Metabolite profiles were examined in leaf samples from two different leaf positions (apical [A] and basal [B] leaves) across three phenological stages (stage 1, stage 2, and stage 3) using targeted metabolomics. A total of 146 metabolites were detected and identified, including 30 organic acids, 24 amino acids, 12 sugars, 27 plant hormones, 18 nucleosides, 27 flavonoids, and 8 other compounds (**Figure 1A, Table S1**). Two-dimensional principal component analysis (PCA) (**Figure 1B**) revealed distinct separations in metabolite profiles across pairwise comparisons (e.g., stages 2A *vs.* 1A, 2B *vs.* 1B, 3A *vs.* 2A, and 3B *vs.*2B), highlighting the significant impact of flush phenology and leaf position along the shoot. Notably, stages 1A *vs.* 1B and 2A *vs.* 2B displayed substantial distribution distances, whereas stages 3A *vs.* 3B showed closer proximities, suggesting stage-specific metabolic differences among leaves in different positions on the same shoots. In three-dimensional PCA (**Figure 1C**), the six sample groups were clearly separated, with PC1, PC2, and PC3 explaining 35.3%, 26.2%, and 9.8% of the total variance, respectively. Volcano plot analysis (*p* < 0.05, fold change > 1.5 or < 1/1.5) identified 96, 84, 76, and 42 differential metabolites (DMs) across stage-wise comparisons (stages 2A *vs.* 1A, 2B *vs.* 1B, 3A *vs.* 2A, and 3B *vs.* 2B) (**Figure 1D-G**). Enrichment analysis using MBRole 2.0 revealed 45, 39, 24, and 19 KEGG pathways (*p* < 0.05), which were visualized as bubble charts (**Figure 1H, Table S2**). Key pathways included plant hormone biosynthesis, metabolic pathways, and starch and sucrose metabolism. Additionally, 67, 77, and 30 DMs were identified for growth position comparisons (stages 1A *vs.* 1B, 2A *vs.* 2B, 3A *vs.* 3B) (**Figure 1I-K**). Pathway enrichment analysis indicated 34, 44, and 19 position-specific metabolic pathways (**Figure 1L, Table S3**), including citrate cycle (TCA cycle), plant hormone biosynthesis, and starch and sucrose metabolism. Among the shared metabolic pathways influenced by phenological stage and growth position, the biosynthesis of plant hormones (map01070) and starch and sucrose metabolism (map00500) garnered our particular interest.

**Figure 1.**
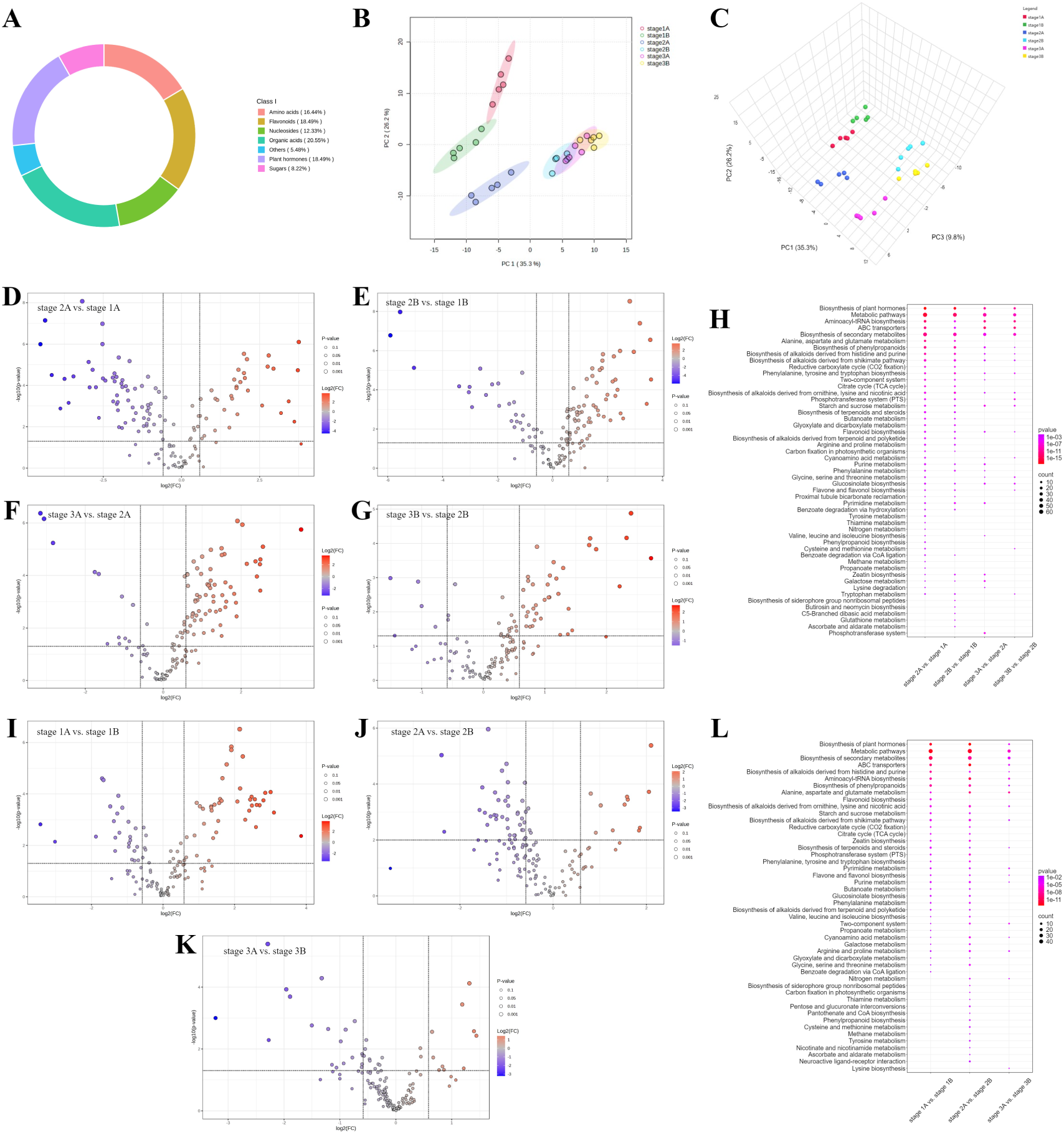
Overview of metabolomics results. Proportions of all metabolite categories (A). The 2D PCA score plot (B). The 3D PCA score plot (C). The DEMs volcano plots for stage 2A *vs.* stage 1A (D), stage 2B *vs.* stage 1B (E), stage 3A *vs.* stage 2A (F), and stage 3B *vs*. stage 2B (G). The combined bubble plot for the metabolic pathway associated with flush phenology (H). The DEMs volcano plots for stage 1A *vs.* stage 1B (I), stage 2A *vs.* stage 2B (J), and stage 3A *vs.* stage 3B (K). The combined bubble plot for the metabolic pathway associated with leaf position (L).

### Proteomics Analysis

Proteomic analysis identified 40,806 peptides and 8,159 proteins, with statistical results detailed in **Figure S1**. The 2D PCA plot (**Figure 2A**) showed clear separations among the six sample groups, with the first two principal components explaining 35.4% and 11.9% of the total variance, respectively. Flush phenology had a stronger impact on protein profiles compared to leaf position. Gene ontology (GO) enrichment analysis (**Figure 2B**) revealed that proteins were primarily involved in metabolic and cellular processes (biological processes, BP), associated with cell parts and cells (cellular components, CC), and linked to catalytic activity and binding (molecular functions, MF). Eukaryotic orthologous group (KOG) analysis highlighted roles in carbohydrate transport and metabolism, energy production and conversion, and amino acid transport and metabolism (**Figure 2C**). Volcano plot analysis identified 3,251, 3,089, 1,517, and 1,131 differentially expressed proteins (DEPs) across stage-wise comparisons (stages 2A *vs.* 1A, 2B *vs.* 1B, 3A *vs.* 2A, and 3B *vs.* 2B) (**Figure 2D-G**). Pathway enrichment revealed that these DEPs were involved in starch and sucrose metabolism, plant hormone signal transduction, zeatin biosynthesis, and carbon metabolism (**Figure 2H, Table S4**). Leaf position comparisons (stages 1A *vs.* 1B, 2A *vs.* 2B, 3A *vs.* 3B) identified 579, 612, and 638 DEPs, respectively (**Figure 2I-K**). Enrichment analysis (**Figure 2L, Table S5**) indicated involvement in photosynthesis, carbon fixation, starch and sucrose metabolism, carbon metabolism, and plant hormone signal transduction. Both proteomic and metabolomic analyses consistently highlighted starch and sucrose metabolism and plant hormone-related signaling pathways as key processes influenced by flush phenology and leaf position.

**Figure 2.**
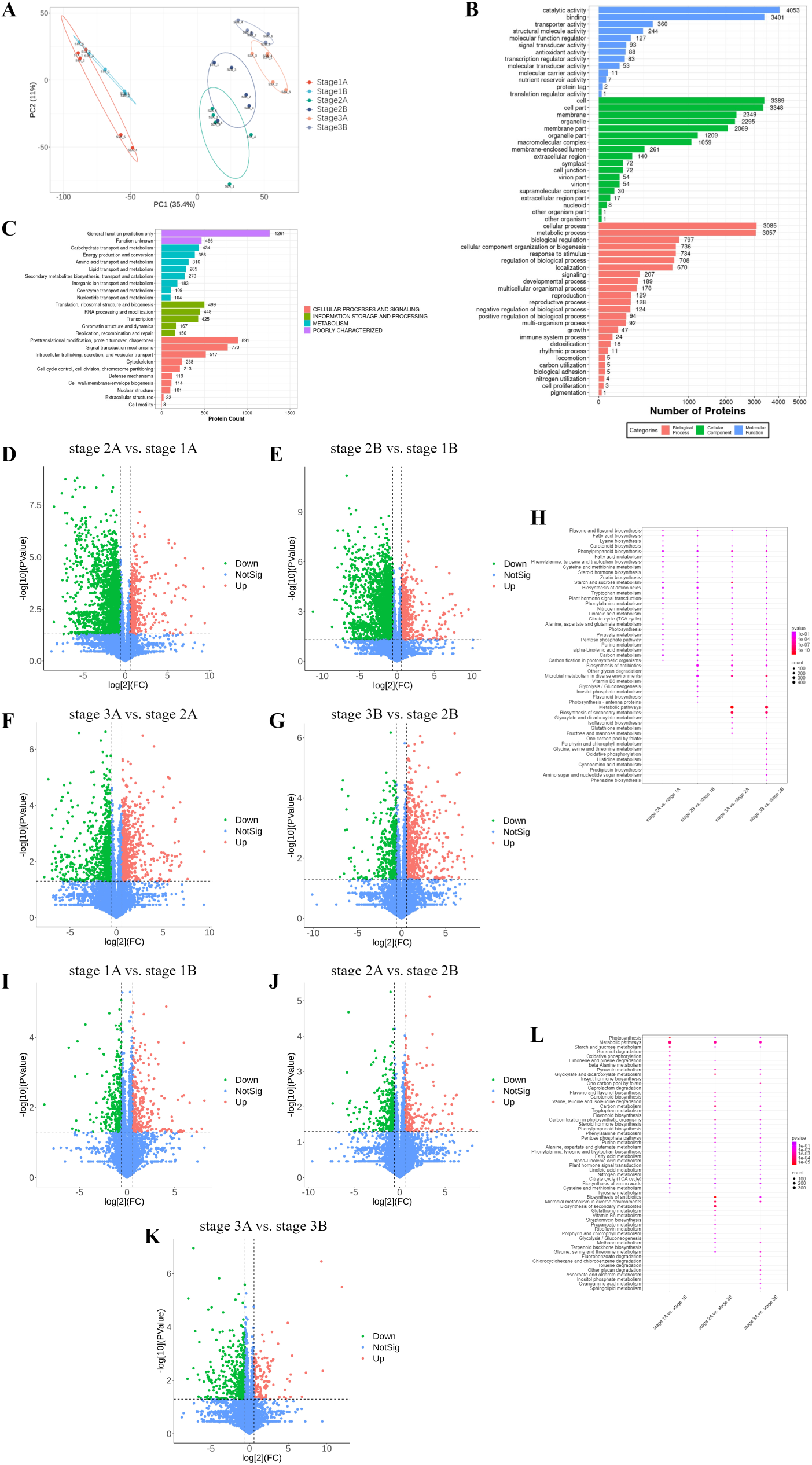
Overview of proteomics results. The 2D PCA score plot (A). GO function annotation (B). KOG analysis (C). The DEPs volcano plots for stage 2A *vs.* stage 1A (D), stage 2B *vs.* stage 1B (E), stage 3A *vs.* stage 2A (F), and stage 3B *vs.* stage 2B (G) The combined bubble plot for the DEP pathway associated with flush phenology (H) The DEPs volcano plots for stage 1A *vs.* stage 1B (I), stage 2A *vs.* stage 2B (J), and stage 3A *vs.* stage 3B (K). The combined bubble plot for the DEP pathway associated with leaf position (L).

### Conjoint Analysis

#### Starch and sucrose metabolism

Metabolomic analysis identified 16 sugar-related metabolites in citrus leaves at various developmental stages, including monosaccharides (e.g., glucose, fructose, fructose-6P, UDP-glucose, glucose-6P, rhamnose, erythritol, xylose, arabitol, sorbitol, and myo-inositol) and polysaccharides (e.g., sucrose, raffinose, trehalose, sucrose-6P, and maltose). From stage 1 to stage 3, monosaccharide levels in both apical and basal leaves, including glucose, fructose-6P, glucose-6P, rhamnose, and xylose, steadily declined (**Figure 3**). In contrast, polysaccharides such as sucrose, maltose, and trehalose accumulated in apical leaves, peaking at stage 2 before decreasing by stage 3, whereas basal leaves exhibited a continuous decline in these polysaccharides from stage 1 to stage 3. Sucrose quantification, performed using a standard curve approach (details in **Supplementary Information and Figure S2**), revealed that in apical leaves, sucrose content reached 11.32 mg·g□ ¹ at stage 2, marking a 73.69% increase from stage 1. However, as development progressed to stage 3, sucrose levels dropped sharply by 71.82%. In basal leaves, sucrose content remained relatively stable between stages 1 and 2 but decreased significantly (50.50%) by stage 3. Notably, at stage 2, sucrose levels in apical leaves were 1.94 times higher than in basal leaves. Vincent et al. (2025) reported a dynamic pattern of starch accumulation in apical and basal leaves, with levels peaking at stage 2 and subsequently declining sharply. By stage 3, starch reserves were nearly exhausted. At stage 2, higher concentrations of starch, sucrose, maltose, trehalose, and raffinose were detected in apical leaves compared to basal leaves, suggesting active mobilization and accumulation of carbohydrate reserves. Notably, the elevated sucrose levels, as the principal metabolite involved in carbon transport, indicate enhanced source-to-sink translocation to support the vigorous metabolic demands of flush development. Six starch-synthesis enzymes including phosphoglucomutase (PGM), ADP-glucose pyrophosphorylase (AGPase), ADP-sugar diphosphatase, starch synthase (SS), granule-bound starch synthase (GBSS) and starch branching enzyme (SBE) were identified. AGPase, the rate-limiting enzyme, peaked in stage 2A, aligning with measured starch content. Starch degradation pathways include α-amylase and β-amylase-mediated hydrolysis to maltose and dextrin, as well as isoamylase (ISA)-catalyzed maltodextrin degradation. From stage 1 to stage 3, α-amylase initially increased before declining, while β-amylase levels gradually decreased. Maltase-glucoamylase degraded maltose to glucose, with its expression levels declining across the stages. Sucrose-6-phosphatase, an enzyme involved in sucrose biosynthesis, catalyzes the formation of sucrose-6P, with higher levels observed in stages 1 and 2. Sucrose is degraded into glucose and fructose by beta-fructofuranosidase (commonly known as invertase, INV), followed by phosphorylation to fructose-6-phosphate via fructokinase (FRK) and hexokinase (HK). The levels of INV, FRK, and HK in both apical and basal leaves steadily declined from stage 1 to stage 3.

**Figure 3.**
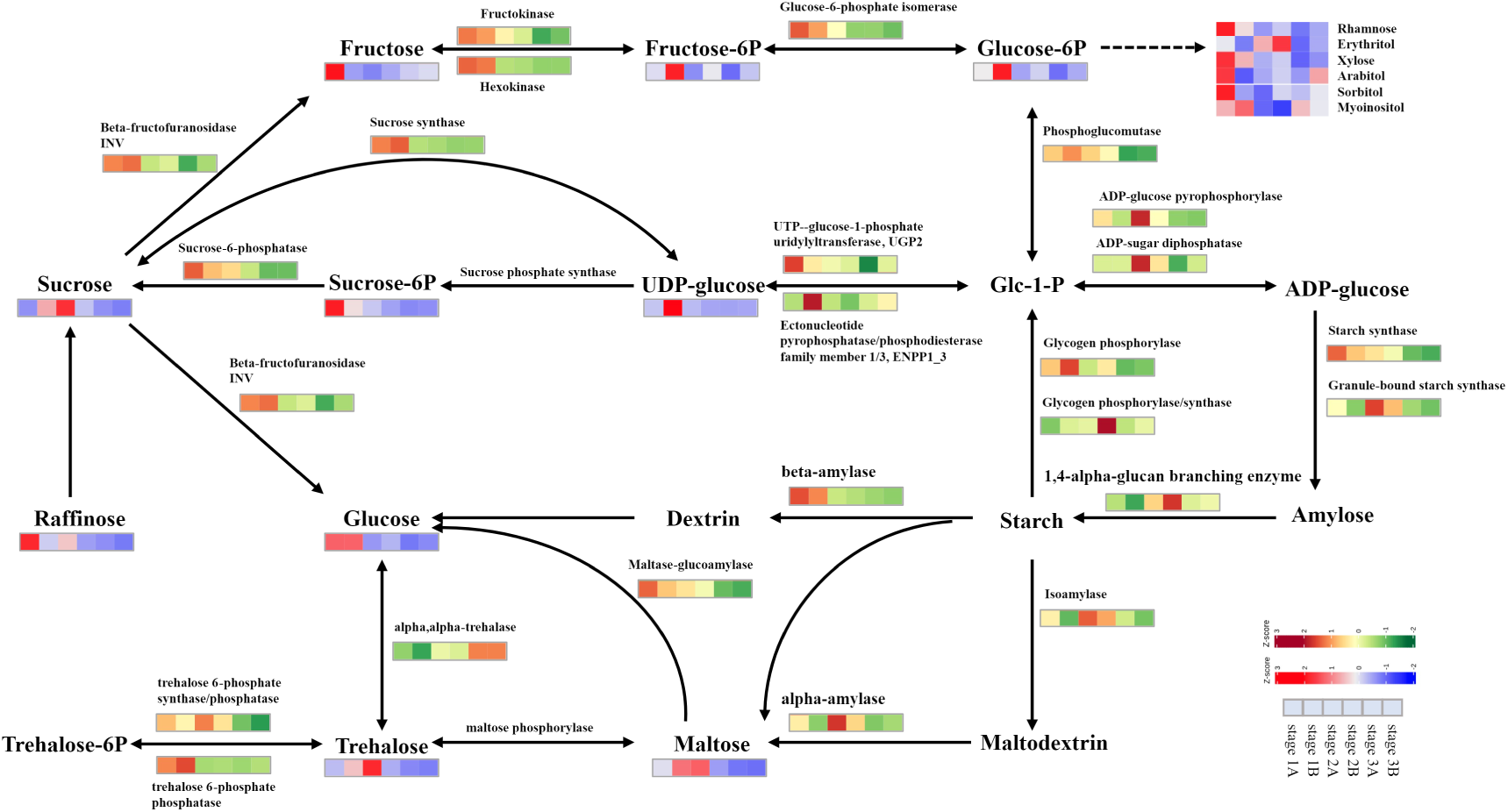
The starch and sucrose metabolic pathway. Metabolites are shown between arrows with a scale of bright red (higher) to blue (lower) showing relative quantities detected by targeted metabolomics, while enzymes involved are shown above or below the arrows with a scale of orange (higher) to purple (lower) showing relative quantities, derived from DIA proteomic analysis.

### Hormone synthesis and signal transduction

#### Auxins

The relative levels of seven auxins, including indole-3-acetamide, indole-3-acetic acid (IAA), indole-3-carboxylic acid, indole-3-acetyl-L-glutamic acid, oxindole-3-acetic acid, indole-3-acetyl-aspartic acid, and indole-3-butyric acid, were determined. Most auxins, except indole-3-butyric acid, were more abundant at stage 3 (**Figure 4A**). Indole-3-acetamide, indole-3-acetic acid, indole-3-acetyl-L-glutamic acid, and indole-3-acetyl-aspartic acid steadily increased from stage 1 to stage 3. Furthermore, indole-3-acetamide, IAA, indole-3-acetyl-aspartic acid, and indole-3-carboxylic acid were more abundant in apical leaves than in basal leaves at stages 2 and 3. IAA, the principal auxin, is synthesized via the tryptophan aminotransferase of Arabidopsis (TAA) /YUCCA pathway and the tryptophan decarboxylation pathway. In the TAA/YUCCA pathway, YUCCA catalyzes the conversion of indole-3-pyruvic acid (IPA) into IAA, with its highest expression at stage 1, followed by a gradual decline. In contrast, amidase, a key enzyme in the tryptophan decarboxylation pathway, converts indole-3-acetamide into IAA. The expression of amidase increased from stage 1 to stage 3, particularly in apical leaves at stage 3, indicating the predominance of this pathway in IAA biosynthesis.

**Figure 4.**
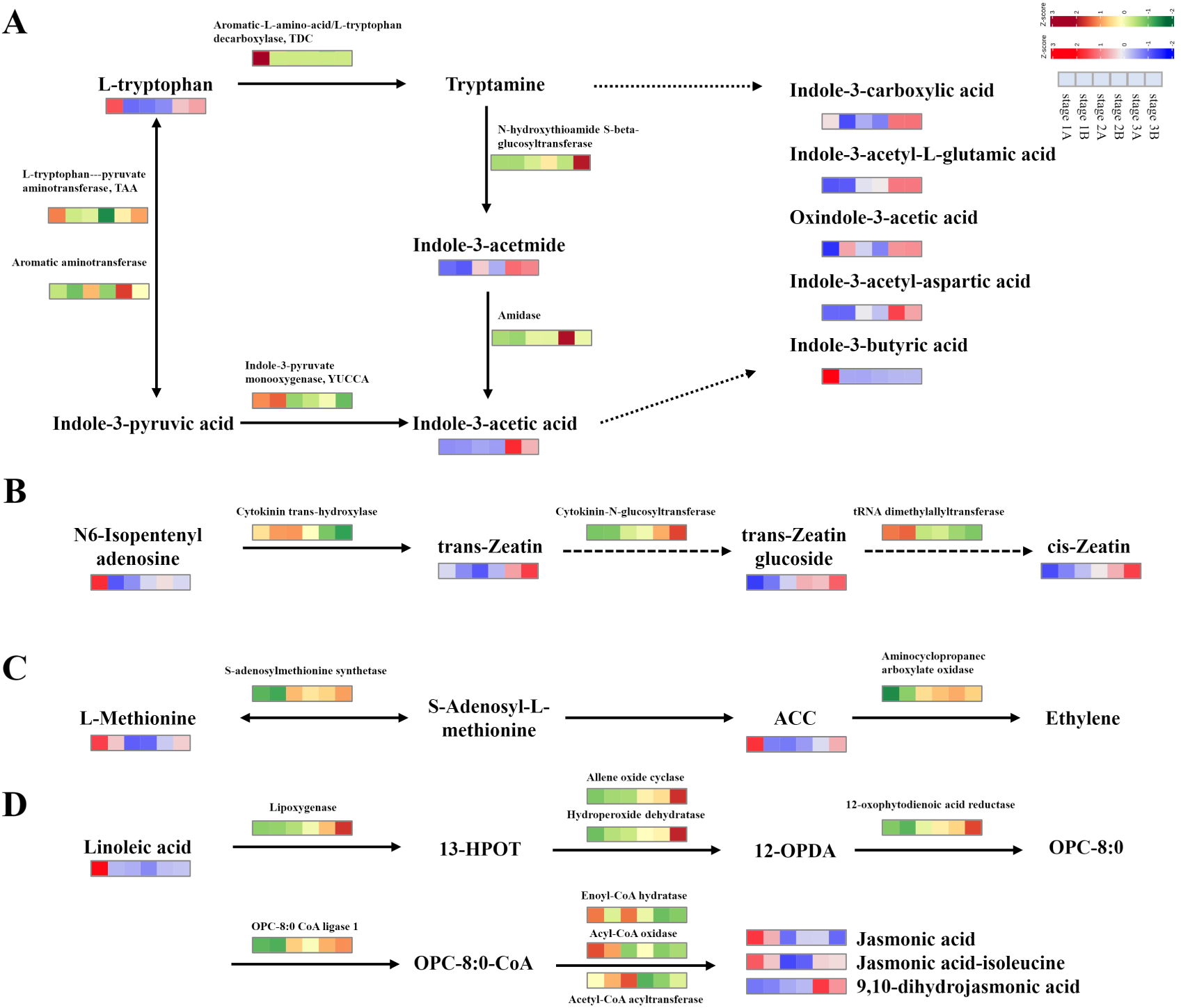
Biosynthesis of plant hormones. The biosynthesis pathway of auxins (A). The biosynthesis pathway of cytokinins (B). The biosynthesis pathway of ethylene (C). The biosynthesis pathway of jasmonic acid (D). Metabolites are shown between arrows with a scale of bright red (higher) to blue (lower) showing relative quantities detected by targeted metabolomics, while enzymes involved are shown above or below the arrows with a scale of orange (higher) to purple (lower) showing relative quantities, derived from DIA proteomic analysis.

Proteomics identified key enzymes in the auxin signal transduction pathway, including auxin resistant 1 (AUX1), transport inhibitor response 1 (TIR1), auxin/indole-3-acetic acid (AUX/IAA), auxin response factor (ARF), small auxin up rna protein (SAUR), and glycine-rich protein 3 (GH3) (**Figure 5A**). AUX1 regulates IAA influx, shaping the auxin gradient. TIR1, the primary auxin receptor, mediates AUX/IAA degradation, releasing ARF to activate downstream gene expression for leaf growth and development. ARF regulates genes linked to auxin responses, while SAUR proteins, downstream of ARF, promote cell wall acidification and expansion, thereby promoting cell growth. GH3 conjugates auxins with amino acids to modulate auxin activity. From stage 1 to stage 3, AUX1, TIR1, and AUX/IAA expression declined, whereas SAUR and GH3 expression increased.

**Figure 5.**
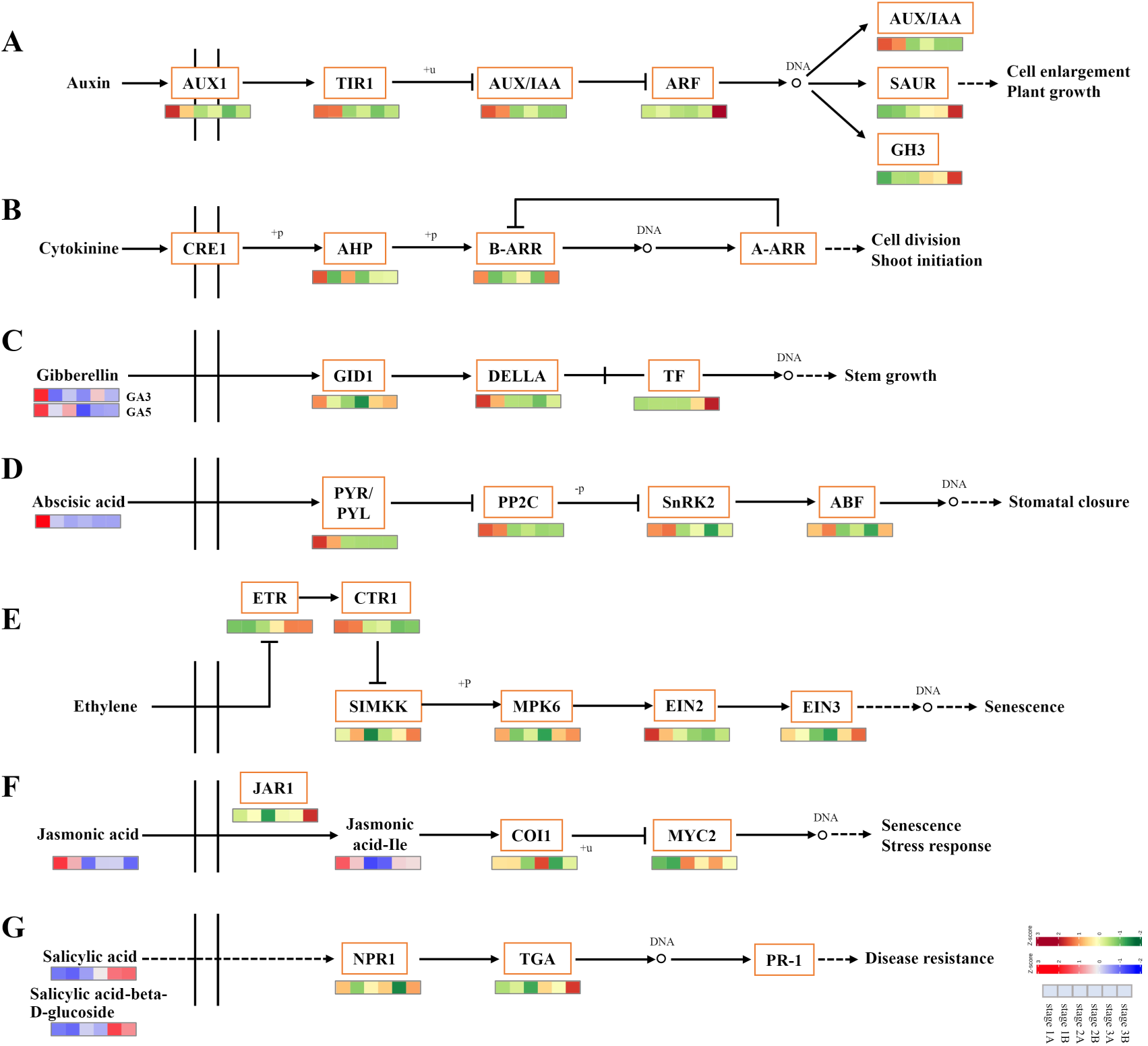
Plant hormone signal transduction pathway. The signal transduction pathway of auxins (A). The signal transduction pathway of cytokinins (B), gibberellin (C), abscisic acid (D), ethylene (E), jasmonic acid (F), and salicylic acid (G). Metabolites are shown between arrows with a scale of bright red (higher) to blue (lower) showing relative quantities detected using targeted metabolomics, while enzymes involved are indicated with a scale of orange (higher) to purple (lower) showing relative quantities, derived from DIA proteomic analysis.

#### Cytokinins (CKs)

Three cytokinins, including trans-zeatin (t-Z), trans-zeatin glucoside, and cis-zeatin, were identified. The levels of t-Z, trans-zeatin glucoside, and cis-zeatin progressively increased from stage 1 to stage 3 (**Figure 4B**). Notably, these cytokinins were more concentrated in basal leaves than in apical leaves during stages 2 and 3. In the zeatin synthesis pathway, three enzymes were identified, including cytokinin trans-hydroxylase, cytokinin-N-glucosyltransferase, and tRNA dimethylallyltransferase. Cytokinin-N-glucosyltransferase, which catalyzes the conversion of trans-zeatin into trans-zeatin glucoside, exhibited increased expression at stages 2 and 3, aligning with the rising levels of trans-zeatin glucoside.

In the cytokinin signal transduction pathway (**Figure 5B**), two proteins were identified: Arabidopsis histidine phosphotransfer protein (AHP) and type-B Arabidopsis response regulator (B-ARR). AHP relays the cytokinin signal to B-ARR, which activates and promotes the expression of genes involved in cell division and expansion, influencing leaf development. From stage 1 to stage 3, AHP and B-ARR levels decreased in apical leaves but increased in basal leaves.

#### Gibberellic Acids

Two forms of gibberellic acid, gibberellic acid 3 (GA_3_) and gibberellic acid 5 (GA_5_), were identified in citrus leaves (**Figure 5C**). GA_3_, recognized as the most active and widely used form, strongly promote cell expansion and carbohydrate remobilization in various plants. In apical leaves, GA_3_ levels initially decreased but rose significantly from stage 1 to stage 3, whereas in basal leaves, GA_3_ levels gradually increased from stage 1 to stage 3. Conversely, GA_5_ levels gradually declined in apical leaves but initially decreased, followed by an increase, in basal leaves. Notably, GA_3_ and GA_5_ concentrations were consistently higher in apical leaves than in basal leaves at all stages. Additionally, in the gibberellin signaling pathway, gibberellin-insensitive dwarfing 1 (GID1) perceives gibberellin signals and facilitates the degradation of DELLA proteins, relieving their inhibitory effects on transcription factors (TFs) and promoting citrus growth and development (Domagalska & Leyser, 2011; Sun, 2010). The observed increase in GA_3_, coupled with higher GID1 and TF levels during stage 3, highlights GA_3_’s critical role in promoting the transition from new bud formation to new shoot development.

#### Abscisic acid

Abscisic acid (ABA) is a key inhibitory plant hormone that regulates bud growth inhibition, dormancy maintenance, and stress responses (Brookbank et al., 2021). As shown in **Figure 5D**, ABA content in citrus leaves was highest at stage 1, followed by a significant decline at stages 2 and 3. Apical leaves exhibited higher ABA levels than basal leaves at this stage, indicating a predominant role in early leaf development. In the ABA signaling pathway, core regulatory components include Pyrabactin resistance/Pyrabactin resistance-like proteins (PYR/PYL), protein phosphatase 2C (PP2C), sucrose non-fermentation-related protein kinase 2 (SnRK2), and abscisic acid response element binding factor (ABF). These proteins were highly expressed during stage 1, indicating an active role for ABA in leaf maturation. During early leaf development, high ABA concentrations suppress unnecessary cell division, maintaining leaf maturity and preventing premature bud activation. As new shoots emerge, ABA levels decline, particularly in apical leaves, facilitating initiation of new bud growth.

#### Ethylene

Ethylene is a well-known regulator of fruit development, ripening, stress responses, and plant growth. Under non-limiting environmental conditions, ethylene typically inhibits shoot growth and development (Reis et al., 2003). In the ethylene biosynthesis pathway, two intermediates, L-methionine and 1-aminocyclopropane-1-carboxylic acid (ACC), were detected. As shown in **Figure 4C**, their levels decreased in stage 2 compared to stage 1. In the ethylene signal transduction pathway, key proteins such as ethylene receptor (ETR), constitutive triple response 1 (CTR1), erine/threonine protein kinase (SIMKK), mitogen-activated protein kinase 6 (MPK6), ethylene-insensitive 2 (EIN2), and ethylene-insensitive 3 (EIN3) were identified. Notably, the expression levels of SIMKK, MPK6, EIN2, and EIN3 were lower in stage 2 (**Figure 5E**), corresponding to the observed reduction in biosynthetic intermediates.

#### Jasmonic acids

Three jasmonic acid (JA) derivatives, including JA, jasmonic acid-isoleucine (JA-Ile), and 9,10-dihydrojasmonic acid, were detected in this study. As shown in **Figure 4D**, from stage 1 to stage 3, 9,10-dihydrojasmonic acid increased gradually, while JA-Ile first decreased and then increased. JA had the highest content in stage 1. JA biosynthesis originates from linoleic acid, which was found in higher concentrations in apical leaves at stage 1. Eight enzymes involved in JA biosynthesis were identified, including lipoxygenase, allene oxide cyclase, hydroperoxide dehydratase, 12-oxophytodienoic acid reductase, OPC-8:0 CoA ligase 1, acyl-CoA oxidase, acetyl-CoA acyltransferase, and enoyl-CoA hydratase. In the JA signaling transduction pathway, three proteins were identified: jasmonic acid resistant 1 (JAR1), coronatine insensitive 1 (COI1), and myelocytomatosis 2 (MYC2). JAR1 catalyzes the conjugation of JA with isoleucine to form JA-Ile, a key bioactive form of JA that influences plant development and stress responses. COI1 binds to JA-Ile and promotes the degradation of downstream transcription factors, while MYC2 is a major transcription factor that regulates the expression of JA-stress-responsive genes. From stage 1 to stage 3, JAR1 first decreased and then increased. COI1 in the top leaves gradually decreased, while COI1 in the bottom leaves initially increased and then decreased. Compared to stage 1, MYC2 content was higher in stages 2 and 3 (**Figure 5F**).

#### Salicylic acids

Two kinds of salicylic acids (Sas), including SA and salicylic acid-beta-D-glucoside (SAG) were detected. SAG serves as a storage form of SA and can be rapidly converted to its active form when needed. From stage 1 to stage 3, the contents of salicylic acid and salicylic acid-beta-D-glucoside gradually increased (**Figure 5G**). In the salicylic acid signaling pathway, two key proteins were identified: non-expressed pathogenesis-related gene 1 (NPR1) and TGACG motif binding factor (TGA). SA, as the most bioactive form, directly activates plant defense genes and induces systemic acquired resistance (SAR), primarily through NPR1-mediated signaling. As SA levels increase, NPR1 becomes activated and translocates to the nucleus, where it stimulates TGA transcription factors to regulate plant growth and immune responses. Interestingly, NPR1 levels gradually decreased in apical leaves but increased in basal leaves from stage 1 to stage 3. Additionally, TGA levels were higher at stage 3 than at stages 1 and 2.

## DISCUSSION

This study advances understanding of citrus flush phenology by integrating metabolomics and proteomics to dissect the coordinated changes in carbohydrate metabolism and hormone dynamics. This work makes four contributions: (1) development of a high-resolution metabolomics and proteomics platform tailored for citrus, (2) elucidation of carbohydrate metabolic dynamics and enzyme regulation during shoot development (a rapid shift in source-sink dynamics), (3) characterization of spatiotemporal hormone signaling pathways involved in bud initiation and growth, and (4) the proposal of a flush-stage integrated metabolic–hormonal regulatory model. These insights provide a comprehensive framework for how flush-forming woody perennials regulate growth and carbon allocation during rapid fluctuations in source-sink balance.

### Multi-omics profiling enhances metabolic annotation accuracy

To achieve high-resolution profiling, we employed a targeted selected ion monitoring (tSIM) metabolomics approach using 121 chemical references. This method ensures high accuracy through reference-based metabolite identification, high sensitivity with a superior signal-to-noise ratio, and strong specificity by minimizing interference. Untargeted metabolomics complemented this strategy by capturing metabolites lacking reference standards, thus expanding metabolic coverage (Schrimpe-Rutledge et al., 2016; Cai et al., 2023). Nevertheless, several sugar phosphate intermediates involved in the Calvin cycle were poorly detected, likely due to their high polarity in chromatography and low ionization efficiency in mass spectrometry (Rende et al., 2019). Future studies may explore derivatization strategies or alternative ionization techniques to improve the detection of these critical metabolites. On the proteomics side, a DIA strategy paired with a spectral library derived from DDA enhanced both protein coverage and quantification precision. DIA allows for comprehensive precursor and fragment ion acquisition, mitigating the limitations of intensity-dependent DDA and reducing missing data (Krasny et al., 2021). This method enhances the depth of protein detection while eliminating the need for labeling, streamlining the workflow. The DDA-generated spectral library served as a high-quality reference, thereby improving the reliability of DIA-based protein quantification. Collectively, this integrated multi-omics approach enabled simultaneous assessment of metabolite pools and enzyme dynamics across key metabolic pathways, improving the resolution and accuracy of metabolic and hormonal network annotation during citrus flush development.

### Flush phenology coordinates sucrose-starch metabolism through enzymatic regulation

Starch and sucrose metabolism play a crucial role in flushing phenology by regulating the source-sink relationship and the balance between carbon fixation and transport. The spatiotemporal expression of key metabolic enzymes serves as a major regulatory factor in this process. During shoot initiation, mature leaves support developing tissues through carbon export and nitrogen remobilization, likely derived in part from Rubisco turnover (Hussain et al., manuscript under revision). Sucrose is a vital photosynthate that serves as the main transported form, while starch acts as the main storage form of carbon fixed during photosynthesis (Stein et al., 2019).

Recent studies have revealed a significant disparity between the rates of carbon fixation and transport capacity in citrus and other trees, a phenomenon known as the fixation-transport capacity difference (Vincent et al., 2025). Sugar transport in plants relies on the phloem, where narrow sieve tubes limit carbon translocation (Rademaker et al., 2017). In citrus, weak carbon transport capacity may lead to excessive carbon fixation with inefficient redistribution, resulting in carbohydrate accumulation in leaves and feedback inhibition of photosynthesis (Welker et al., 2022; Li & Vincent, 2022; Li et al., 2024). Moreover, despite their high photosynthetic potential (maximum rate of carboxylation or of RuBP regeneration), citrus leaves exhibit low stomatal conductance, restricting CO□ uptake well below its theoretical maximum (Hussain et al., 2024). Enhancing carbon transport efficiency could alleviate photosynthetic inhibition and ultimately improve crop productivity (Li et al., 2025). Our results show that sucrose, trehalose, and maltose concentrations in apical leaves peak at stage 2 and decline markedly by stage 3 during flush phenology. This pattern suggests not only the progressive depletion of carbohydrate reserves in stage 3 but also a reduced availability of metabolically active monosaccharides within the mature leaves. Hussain et al. (under review) have demonstrated that photosynthesis begins to decline at stage 2 due to reduced stomatal conductance, and further declines in stage 3 as carboxylation decreases and Rubisco content declines. The decline of photosynthetic capacity reported by Hussain et al. (under review) is supported here by the shift to starch metabolism and declining concentrations of photosynthates, as well as by the shifts in amino acid metabolism and Krebbs-cycle activity (**Figure S3**). Given that total transport activity is limited (Vincent et al., 2025), the strategy of temporarily retooling metabolism to export available photosynthates, and remobilize amino acids contained in Rubisco, may enhance the total photosynthetic capacity by allowing new leaves to initiate photosynthesis quickly. Indeed, by the time citrus leaves are fully expanded, before they reach mature chlorophyll content, they are already net exporters of photosynthates (Goldschmidt & Koch, 2017). Consequently, mature apical leaves may undergo substantial functional suppression as carbon and nitrogen resources are remobilized to support the growth of newly emerging leaves. This pattern is closely associated with the expression of key starch-metabolizing enzymes, including AGPase, α-amylase, and isoamylase. AGPase, the rate-limiting enzyme in starch biosynthesis, reaches peak expression at stage 2, indicating an upregulation of carbohydrate synthesis (Wang et al., 2023). During this phase, the photosynthetic capacity of citrus branches exceeds their capacity to transport carbon, leading to substantial accumulation of starch and sucrose, particularly in apical leaves (Vincent et al., 2025). This suggests that citrus preferentially stores carbohydrates in regions destined for future shoot elongation (apical stem and leaves), ensuring sufficient carbon availability for rapid growth. At stage 3, the levels of sucrose, starch, trehalose, and maltose in both apical and basal leaves decrease significantly, with starch being nearly depleted, and even sucrose, which is usually stable to regulate phloem loading and osmotic potentials declining strongly. This decline may be attributed to the combination of reduced photosynthetic capacity and increased carbon remobilization during new shoot formation (Vincent et al, 2025; Hussain et al, under review). Together, these carbon and nitrogen metabolism results support the strong shift found by Hussain et al. (*under review*) from mature leaves producing and storing carbohydrates to supplying both carbohydrates and nitrogen to new leaves, at the expense of photosynthesis in the mature leaves.

### Hormonal regulation orchestrates bud emergence and new shoot development

Hormonal regulation plays a crucial role in bud formation, growth, and shoot development. Our study demonstrates dynamic changes in auxins, cytokinins, gibberellins, abscisic acid, jasmonic acid, and salicylic acid, highlighting their coordinated roles in regulating citrus flush phenology. According to the polar transport hypothesis of auxin, auxin’s directional movement within the stem is essential for shoot formation and growth. IAA, as a key regulator of plant growth, plays a vital role in cell expansion and differentiation (Teale et al., 2006). Our study observed a gradual increase in the levels of IAA and its derivatives—indole-3-acetamide, indole-3-acetyl-L-glutamate, and indole-3-acetyl-aspartate—during the formation of shoot buds and subsequent shoot development. This increase was accompanied by the upregulation of IAA biosynthesis-related enzymes, including N-hydroxythioamido-S-β-glucosyltransferase and amidase, as well as signal transduction proteins such as ARF and GH3. These findings highlight the central role of IAA from mature leaves in promoting cell expansion and differentiation occurring in the new shoots, further supporting the concept of auxin polar transport, as IAA levels were consistently higher in apical leaves than in basal leaves.

Cytokinin profiling revealed an increase in trans-zeatin glucoside and cis-zeatin levels from stage 1 to stage 3, whereas t-Z, the most bioactive cytokinin, peaked at stage 3. CKs promote bud activation via cell division, but once bud development is initiated, auxins transported from the growing shoot apex may suppress further cytokinin synthesis (Tanaka et al., 2006; Hussain et al., 2021). In perennial species, t-Z is continuously synthesized in the root system and transported to aerial tissues via the xylem, establishing a long-distance signaling axis that plays a critical role in coordinating shoot regeneration and spatial distribution throughout different developmental phases or in response to environmental cues. The elevated t-Z levels observed at stage 3 likely contribute to the maintenance of stem cell identity within the shoot apical meristem and promote sustained cell proliferation. This hormonal environment supports longitudinal shoot elongation and facilitates the establishment of primary tissue architecture in newly emerging shoots.

GA□ showed a pronounced increase in apical leaves from stage 2 to stage 3, accompanied by the upregulation of GID1 and downstream signaling factors, suggesting its key role in promoting the transition from bud initiation to shoot outgrowth (Domagalska & Leyser, 2011; Sun, 2010). Beyond its classical function in promoting cell division and elongation, GA□ also modulates carbon allocation by enhancing starch degradation and increasing sucrose availability to support shoot growth (Kim et al., 2006; Nanda et al., 1968; Singh & Chandra, 2021). Moreover, GA□ may influence the expression of key enzymes and transporters involved in sucrose synthesis (SPS, SUSY) and phloem loading (SUTs, SWEETs), facilitating efficient carbon allocation to the growing sink (Palmai 2025; Rossetto et al., 2003; Murcia et al., 2016). ABA levels were highest at stage 1 but significantly declined at later stages. Besides, the high expression of ABA signaling components (PYR/PYL, PP2C, SnRK2, and ABF) at stage 1 reinforces ABA’s role in inhibiting premature bud activation and maintaining dormancy (Pan et al., 2021).

Ethylene biosynthesis intermediates (L-methionine and ACC) and key signaling proteins (EIN2, EIN3, and MPK6) decreased from stage 1 to stage 2, suggesting that ethylene downregulation contributes to bud initiation. Since JA and JA-Ile are known to suppress bud development by inhibiting cell division (Wasternack & Hause, 2013), the decline in their levels during stage 2 likely promotes bud development. Besides, JAs have been recognized as key regulators of inducible defense responses, playing a crucial role in protecting plants against herbivorous insects and microbial pathogens (Marquis et al., 2020; Li et al., 2022). Notably, the increase in JA-Ile from stage 2 to stage 3 coincides with new shoot formation and growth, suggesting its role in regulating both growth and defense. Both SA and its storage form, salicylic acid-beta-D-glucoside (SAG), showed a steady increase from stage 1 to stage 3. Given SA’s well-established role in systemic acquired resistance (Mishra et al., 2024; Monte, 2023), its increasing trend may reflect a defense priming mechanism as young leaves emerge. Together, these signals of increasing auxins, GAs, and cytokinins drive the mature leaf shift in carbon and nitrogen metabolism to support new shoot growth.

### An integrated model of metabolic and hormonal regulation during flushing

This study integrated metabolomics and proteomics to elucidate the intricate regulatory networks governing carbon metabolism and hormonal dynamics during citrus flush phenology. A comprehensive model (**Figure 6**) is proposed to illustrate the spatiotemporal regulation of carbohydrate metabolism and endogenous hormone signaling across distinct flush phenological stages. From stage 1 to stage 2, leaves—particularly apical leaves—enter a carbon storage phase. On one hand, apical leaves initiated signaling for bud outgrowth, characterized by coordinated hormonal regulation involving IAA, trans-zeatin (t-Z), GA□, and ABA, along with their respective signaling enzymes. Notably, significant increases in IAA levels and the expression of signaling enzymes, such as CH3 and SAUR, were observed in both apical and basal leaves. On the other hand, apical leaves exhibited marked starch accumulation. This is likely because they compete less successfully than basal leaves for export, but this has the result of locally available carbon reserves for bud activation (Vincent et al., 2025). This was supported by the upregulation of key enzymes involved in starch metabolism, including AGPase, α-amylase, and isoamylase. Carbon export from basal leaves increased significantly at stage 2 (Vincent et al., 2025), which may partially explain the lower starch content in basal leaves compared to apical ones.

**Figure 6.**
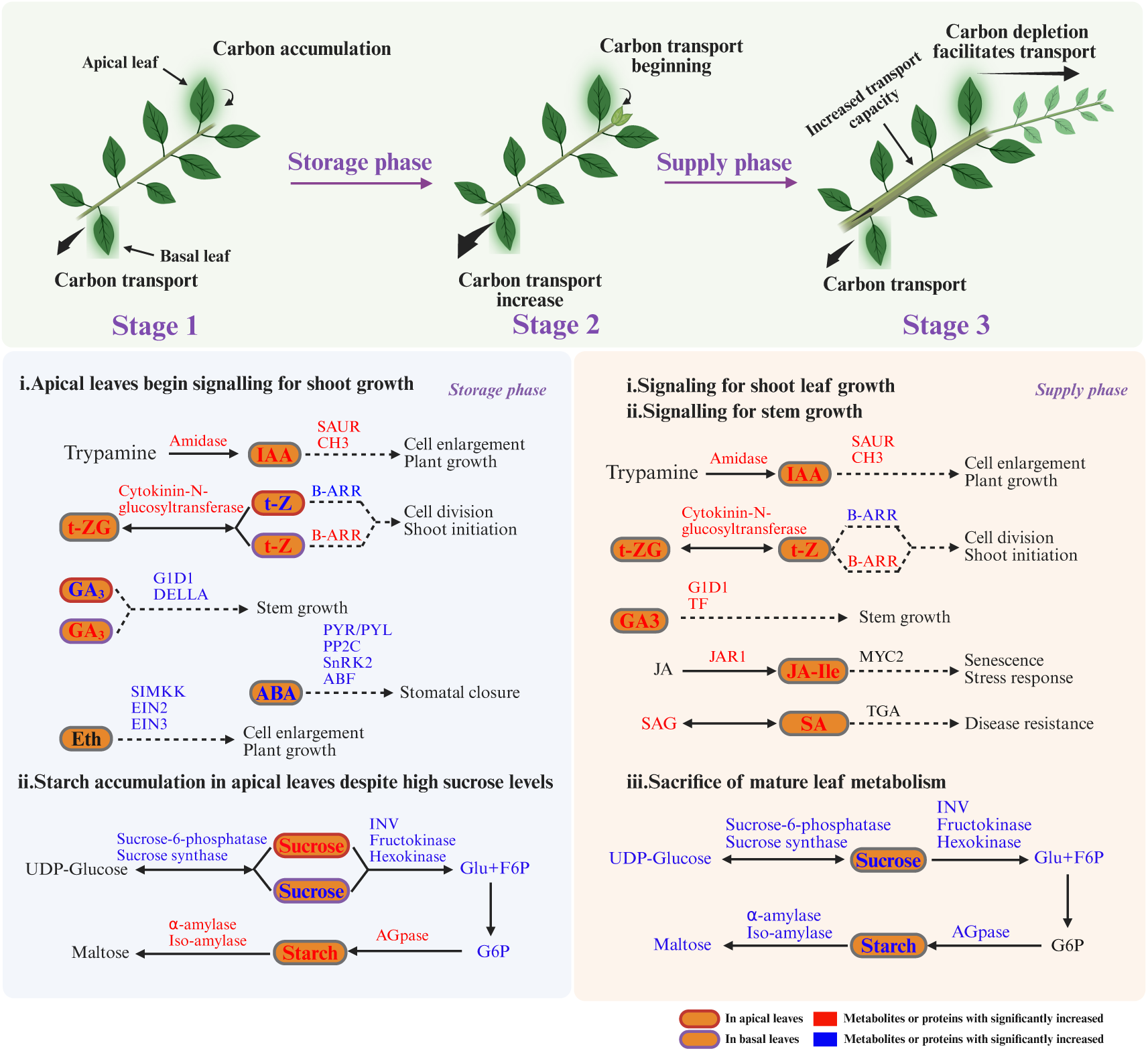
A flush-stage integrated model of metabolic and hormonal regulation.

During the transition from stage 2 to stage 3, both apical and basal leaves shift into a carbon-export phase to sustain active shoot development. Compared with stage 2, stage 3 leaves showed elevated levels of IAA, t-Z, GA□, JA-Ile, and SA, as well as upregulation of associated signaling enzymes such as CH3, SAUR, and CID1, collectively promoting shoot elongation and thickening. Consistent with patterns observed in other woody perennials, citrus stems or branches undergo continuous radial growth mediated by vascular cambium activity, leading to the progressive accumulation of secondary xylem and phloem tissues (Evert, 2006; Fischer et al., 2019). The gradual development of phloem enhances carbohydrate transport capacity and plays a pivotal role in supporting vigorous shoot growth (Ray and Savage, 2021; Heo et al., 2014). Concurrently, mature leaves exhibited reduced metabolic activity, as indicated by significant declines in starch and sucrose content, alongside downregulation of associated metabolic enzymes including AGPase, α-amylase, sucrose-6-phosphatase, INV, and fructokinase. Enhanced carbon transport capacity in the developing stem or branch further facilitates the redistribution of assimilates to rapidly expanding shoot tissues.

Collectively, this study establishes a comprehensive molecular framework elucidating the integration of carbohydrate metabolism and hormone signaling throughout citrus flush development. The proposed flush-stage integrated model of metabolic and hormonal regulation offers mechanistic insights into stage-specific physiological transitions, providing a theoretical basis for precision regulation of shoot growth and carbon allocation. These findings hold significant potential for improving citrus cultivation strategies and enhancing overall productivity.

## Acknowledgments

We thank Myrtho Pierre at the University of Florida for her assistance in the daily management of experimental plants.

## Supplementary Data

**Table S1** Information of metabolites and their analytical conditions.

**Table S2** The obtained kegg pathways associated with phenological stages by metabolomics.

**Table S3** The obtained kegg pathways associated with leaf position by metabolomics.

**Table S4** The obtained kegg pathways associated with phenological stages by proteomics.

**Table S5** The obtained kegg pathways associated with leaf position by proteomics.

**Figure S1** The number of identified proteins in each group.

**Figure S2** Comparison of sucrose content in apical and basal leaves at three phenological stages (n = 5). Data presented as mean ± SEM; \**p* < 0.05, \*\**p* < 0.01, \*\*\**p* < 0.001.

## Author contributions

KC, SBH, XL, QM: Investigation; KC: Data Curation, Visualization, Writing-Original Draft; CV and YW: Conceptualization, Methodology, Project Administration, Supervision, Writing-Review & Editing.

## Data availability statement

The data that support the findings of this study are openly available in Zenodo platform at https://zenodo.org, reference number [16790344].

## Abbreviations

ABA: Abscisic acid
ABF: Abscisic acid response element-binding factor
ACC: Aminocyclopropane-1-carboxylic acid
AHP: Arabidopsis histidine phosphotransfer protein
AGPase: ADP-glucose pyrophosphorylase
ARF: Auxin response factor
AUX1: Auxin influx carrier
AUX/IAA: Auxin/Indole-3-acetic acid proteins
B-ARR: Type-B Arabidopsis response regulator
INV: beta-fructofuranosidase (commonly known as invertase)
BRC1: Branched 1
CKs: Cytokinins
COG: Clusters of orthologous groups
COI1: Coronatine insensitive 1
CTR1: Constitutive triple response 1
DDA: Differentially detected analytes
DELLA: DELLA proteins (repressors of gibberellin signaling)
DMs: Differential metabolites
DIA: Data-independent acquisition
EIN2: Ethylene-insensitive 2
EIN3: Ethylene-insensitive 3
ETR: Ethylene receptor
GA3: Gibberellin A3
GA5: Gibberellin A5
GAs: Gibberellins
GBSS: Granule-bound starch synthase
GH3: Gretchen Hagen 3
GID1: Gibberellin-insensitive dwarfing 1
GO: Gene ontology
HB21: Homeobox 21
HB40: Homeobox 40
HB53: Homeobox 53
IAA: Indole-3-acetic acid
JA-Ile: Jasmonoyl-isoleucine
JAR1: Jasmonic acid resistant 1
MPK6: Mitogen-activated protein kinase 6
MYC2: Myelocytomatosis 2
NCED3: 9-cis-epoxycarotenoid dioxygenase 3
NPR1: Non-expressed pathogenesis-related gene 1
PCA: Principal component analysis
PGM: Phosphoglucomutase
PP2C: Protein phosphatase 2C
PYR/PYL: Pyrabactin resistance/Pyrabactin resistance-like proteins
QC: Quality control
SAG: Salicylic acid-beta-D-glucoside
SA: Salicylic acid
SAUR: Small auxin-up RNA proteins
SBE: Starch branching enzyme
SIMKK: Erine/threonine protein kinase
SnRK2: Sucrose non-fermentation-related protein kinase 2
SS: Soluble starch synthase
TAA/YUCCA: Tryptophan aminotransferase of Arabidopsis/Auxin biosynthesis pathway components
TEAB: Triethylammonium bicarbonate
TGA: TGACG motif binding factor
TIR1: Transport inhibitor response 1
t-Z: trans-Zeatin

